# GluK2 is a target for gene therapy in drug-resistant Temporal Lobe Epilepsy

**DOI:** 10.1101/2023.04.13.536748

**Authors:** Céline Boileau, Severine Deforges, Angélique Peret, Didier Scavarda, Fabrice Bartolomei, April Giles, Nicolas Partouche, Justine Gautron, Julio Viotti, Haley Janowitz, Guillaume Penchet, Cécile Marchal, Stanislas Lagarde, Agnès Trebuchon, Nathalie Villeneuve, Julie Rumi, Thomas Marissal, Roustem Khazipov, Ilgam Khalilov, Fanny Martineau, Marine Maréchal, Anne Lepine, Mathieu Milh, Dominique Figarella-Branger, Etienne Dougy, Soutsakhone Tong, Romain Appay, Stéphane Baudouin, Andrew Mercer, Jared B. Smith, Olivier Danos, Richard Porter, Christophe Mulle, Valérie Crépel

## Abstract

**Objective:** Temporal lobe epilepsy (TLE) is characterized by recurrent seizures generated in the limbic system, particularly in the hippocampus. In TLE, recurrent mossy fiber sprouting from dentate gyrus granule cells (DGCs) creates an aberrant epileptogenic network between DGCs which operates via ectopically expressed GluK2/GluK5-containing kainate receptors (KARs). TLE patients are often resistant to anti-seizure medications and suffer significant comorbidities; hence there is an urgent need for novel therapies. Previously we have shown that GluK2 knockout mice are protected from seizures. This study aims at providing evidence that downregulating KARs in the hippocampus using gene therapy reduces chronic epileptic discharges in TLE.

**Methods:** We combined molecular biology and electrophysiology in rodent models of TLE and in hippocampal slices surgically resected from patients with drug-resistant TLE.

**Results:** Here we confirmed the translational potential of KAR suppression using a non-selective KAR antagonist that markedly attenuated Interictal-like Epileptiform Discharges (IEDs) in TLE patient-derived hippocampal slices. An adeno-associated virus (AAV) serotype-9 vector expressing anti-*grik2* miRNA was designed to specifically downregulate GluK2 expression. Direct delivery of AAV9-anti *grik2* miRNA into the hippocampus of TLE mice led to a marked reduction in seizure activity. Transduction of TLE patient hippocampal slices reduced levels of GluK2 protein and, most importantly, significantly reduced IEDs.

**Interpretation:** Our gene silencing strategy to knock down aberrant GluK2 expression demonstrates inhibition of chronic seizure in a mouse TLE model and IEDs in cultured slices derived from TLE patients. These results provide proof-of-concept for a gene therapy approach targeting GluK2 KARs for drug-resistant TLE patients.

## Introduction

Epilepsy affects up to 1% of people from Western countries^1^. Temporal lobe epilepsy (TLE) is the most prevalent type of focal epilepsy in adults. TLE is characterized by the occurrence of unpredictable and recurrent seizures, which are generated in the medial temporal lobe of the brain, notably in the hippocampus^2^. Recurrent seizures affect normal brain function leading to diminished cognitive capabilities, including progressive memory loss and emotional deficits^3–5^. Patients with epilepsy may also face social stigmatization and isolation, have reduced employment opportunities, and fail to reach educational targets^6, 7^. The World Health Organization estimates that the risk of premature death in individuals with epilepsy is three times higher than that of the general population^8^.

A substantial proportion of epilepsy patients (estimates vary between 30-40%) are resistant to classical drug therapy^1, 9^. In drug-resistant TLE, the surgical resection of the epileptic focus is a last resort intervention. However, fewer than 1% of patients are referred for surgery^10^ because of the potential side effects of removing brain tissue^11^ and seizure freedom is not achieved in all resected patients^12^. Numerous new anti-seizure medications have been developed; however, the proportion of patients with drug-resistant TLE has not been markedly reduced^9, 13^. Clearly, novel therapeutic strategies for the treatment of drug-resistant TLE are urgently required to address the unmet needs of these patients.

In the hippocampal circuit of TLE patients (and in TLE disease models), a fraction of excitatory and inhibitory neurons degenerate, and surviving neurons can form new aberrant synaptic connections^14–16^. Synaptic reorganization in TLE is well documented in the dentate gyrus (DG). Mossy fibers that project from the DG cells (DGCs) to CA3 pyramidal cells sprout recurrent axons to form aberrant glutamatergic excitatory synapses between DGCs, in animal TLE models and in patients^17^. These abnormal synapses operate via ectopically expressed kainate receptors (KARs)^18^ which increase the excitability of the DG^19^.

We have previously shown that pharmacological inhibition of KARs or genetic deletion of GluK2-KARs reduces Interictal-like Epileptiform Discharges (IEDs) and chronic seizures in mouse epileptic hippocampal slices and in a pilocarpine mouse model of TLE, indicating that KARs play a prominent role in the generation of chronic seizures ^20, 21^. GluK2-containing KARs, therefore, represent a rational target for novel therapeutic strategies to treat TLE patients.

Here we show that the pharmacological inhibition of KARs can alleviate IEDs in acute hippocampal slices from TLE patients. Based on this proof-of-concept study, an AAV9 (adeno-associated virus serotype-9) gene therapy vector was designed to deliver small non-coding miRNAs that downregulate GluK2 expression in the human hippocampus via the translational repression of *grik2* mRNA. We demonstrate that hippocampal delivery of AAV9-anti *grik2* miRNA markedly reduces chronic seizure activity in TLE mice and IEDs in cultured slices derived from TLE patients. These results support the use of gene therapy targeting GluK2-containing KARs for the treatment of drug-resistant TLE patients.

## Materials and Methods

### Ethics

*In vivo* mouse experiments were performed in accordance with the European community council directives (2010/63/UE) and received approval from the French Ministry for Research, after ethical evaluation by the institutional animal care and use committee of Aix-Marseille University (protocol number: #9896-201605301121497v11). For experiments using human tissues, all patients gave their written consent and protocols were approved: for AP-HM by Inserm (N° 2017-00031), AP-HM (N° M17-06) under the supervision of CRB TBM/AP-HM (AC: 2018-3105)^22^; and for CHU Bordeaux by CNRS (Codecoh DC-2020-3863).

### Design and production of viral vectors targeting GluK2 by RNA interference

RNAi sequences were designed using the Smart selection protocol^23^. We compared the efficiency of 3 RNAi sequences (RNA#1: CATAGCTAATGCCTGTTTT, RNA#2: TGACATAGAAGGAATCTTT, RNA#3: AGACTTGGGTATTTTCTGT) expressed as shRNAs under the control of the mouse U6 promoter in a lentiviral vector (LV) (Table 1). To achieve effective knock-down, the 19mer shRNA#1 was converted into a 22mer using a human miR30 structure to create miRNA#1^24^. LV or AAV9 vectors were further designed to express the miRNA#1 under the human synapsin 1 (hSYN1) promoter (Table 1). The following LV or AAV9 constructs were used as controls for cultured neurons and mouse organotypic slices: LV.hSYN1.GFP (LV-GFP) or LV.hSYN1.TdTomato.U6.shScramble (LV-shScramble); AAV9.hSYN1.GFP (AAV9-GFP) or AAV9.hSYN1.GFP.miR30.scramble (AAV9-miRScramble) (Fig 2)(Table 1). In chronic epileptic mice and in human organotypic slices, AAV9-miRScramble or AAV9-GFP were used as controls (referred to as AAV9-control in Fig 4, 5). LV-shScramble and AAV9-miRScramble expressed the scramble RNAi sequence TTTGTGAGGGTCTGGTC and TAATGTTAGTCATGTCCACCGCT, respectively. LV vector batches were designed and produced at Univ. Bordeaux (France). AAV9 vectors were provided by Regenxbio (Rockville, USA).

**Figure 1:**
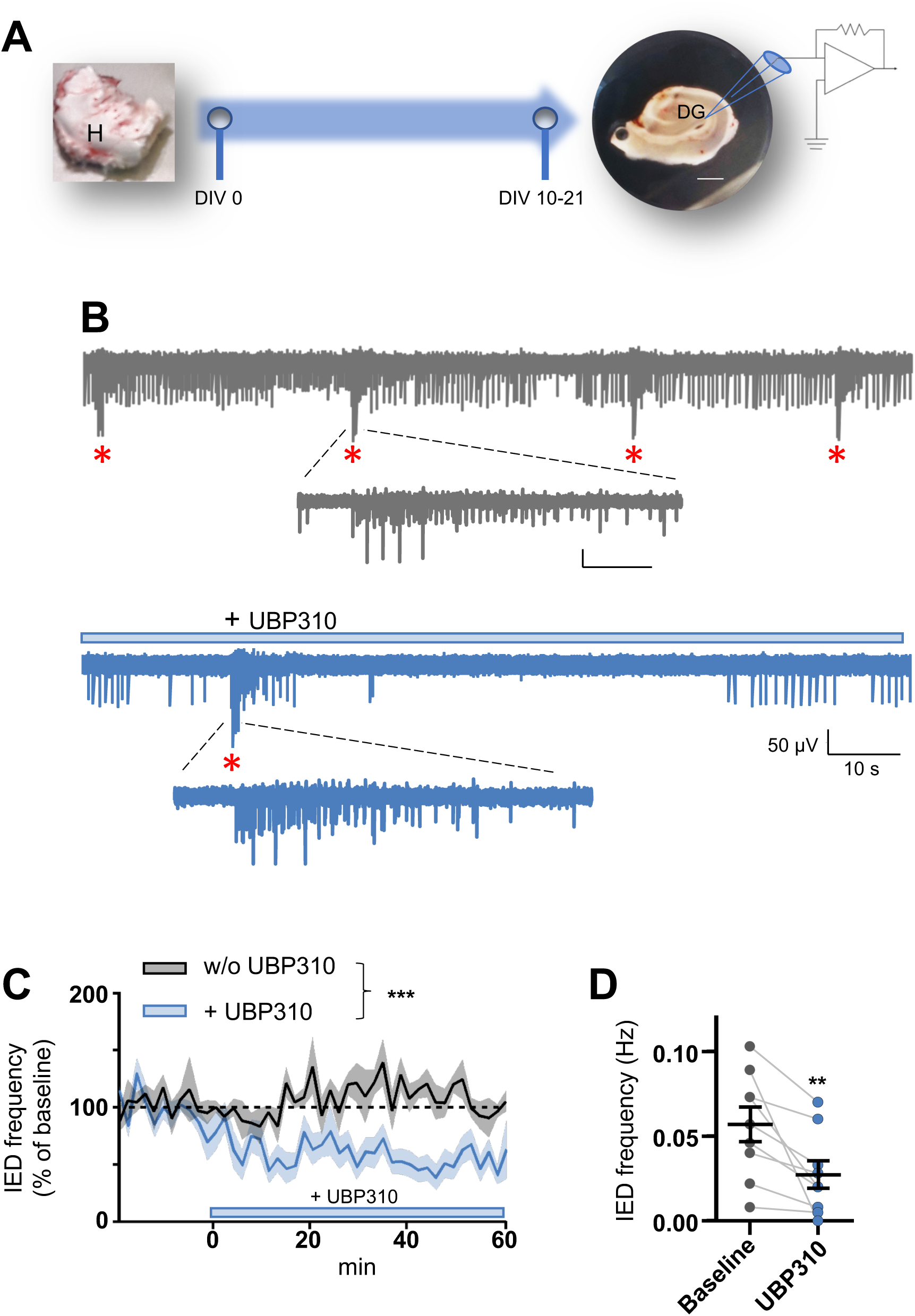
Pharmacological inhibition of KARs reduces the frequency of interictal-like epileptiform discharges in human epileptic hippocampus. **(A)** Illustration of the experimental design and timeline for human organotypic hippocampal slices and LFP (local field potential) recordings including examples of a surgically resected hippocampus (H, left) from a patient with temporal lobe epilepsy (TLE) before slicing and an organotypic slice (right) of the resected hippocampus (scale bar = 5 mm). **(B)** Representative traces of LFPs recorded in a hippocampal organotypic slice derived from surgical resection, before (top) and after (bottom) UBP310 application; enlarged Interictal-like Epileptiform Discharges (IEDs) are shown in the right panel. **(C)** Frequency of IEDs as a function of time in control conditions (grey line, without UBP310), and before and after UBP310 application (blue line, n=9 slices from 4 patients) (***, p=0.0003 by two-way ANOVA, F(1, 14)=22.03). The “0” in the x-axis represents the time point when UBP310 was applied to the recording bath and data were normalized to the average value of the baseline (i.e., the 15 first minutes of recording in control, and the 15 minutes prior to UBP310 application). **(D)** Frequency of IEDs measured in organotypic slices for 15 min before and 40-45 min after UBP310 application (n=9 slices from 4 patients, **, p=0.0039 by Wilcoxon matched-pairs signed rank test). In this and following figures, data are represented as mean ± SEM.

**Figure 2:**
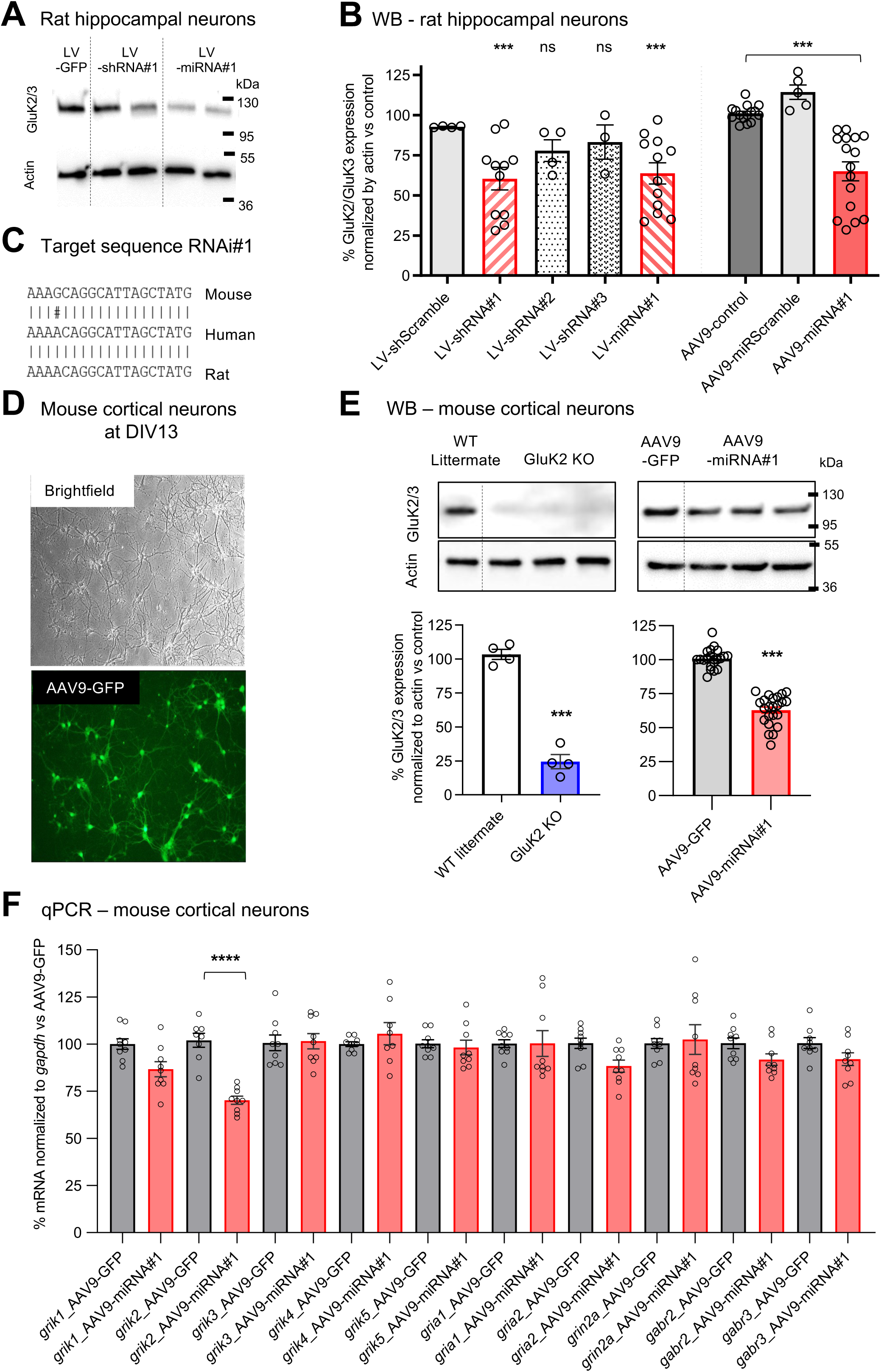
Design and validation of GluK2 knock-down by RNA silencing. **(A)** Western blot of GluK2/GluK3 protein and β actin obtained from rat hippocampal neurons in culture following transduction of lentiviral vectors (LV) to knock-down *grik2* with either an shRNA or an miRNA construct (sequence RNAi#1). **(B)** Graph representing the relative levels of proteins labeled with a GluK2/3 antibody normalized to β actin vs the value in control conditions for hippocampal neurons transduced with the various viral constructs tested (***, p< 0.001 by 1-way ANOVA and Dunnett’s multiple comparisons tests vs LV-GFP for LV vectors and vs AAV9-GFP for AAV9 vectors). **(C)** Complementary RNA sequence (sequence RNAi#1) used to knock-down *grik2* in rodents and human. **(D)** Mouse cortical neurons in culture transduced with an AAV9-hSYN1-eGFP viral construct (AAV9-GFP). Direct eGFP fluorescence was observed in 89±2% of DIV13 neurons. **(E)** Representative western blotting of proteins labeled with the GluK2/3 antibody and actin for mouse cortical neurons of GluK2 KO mice vs wild-type controls (dot plot and mean ± SEM; ***, p<0.001 by unpaired t test) and wild-type mice transduced with AAV9 viral vectors expressing the miRNA (sequence RNAi#h1) under the control of the hSYN1 promoter to knock-down *grik2* (n=8 independent neurons cultures) vs cultures transduced with a control AAV9 vector. **(F)** Comparative transcriptomic analyses by qPCR of mouse cortical neurons transduced with AAV9 viral vectors expressing the miRNA (sequence RNAi#h1). Graph representing the relative levels of Glutamate receptor sub-units (Kainate: *grik1* to *grik5*; AMPA: *gria1* and *gria2*; NMDA: *grin2a*) and GABA_A_ receptor subunits (*gabr2* and *gabr3*) mRNA normalized to *gapdh* vs the value in AAV9-GFP control conditions (dot plot and mean ± SEM; n=3 independent neurons culture; ***, p<0.001 by one-way ANOVA and Dunnett’s multiple comparisons test).

**Figure 3:**
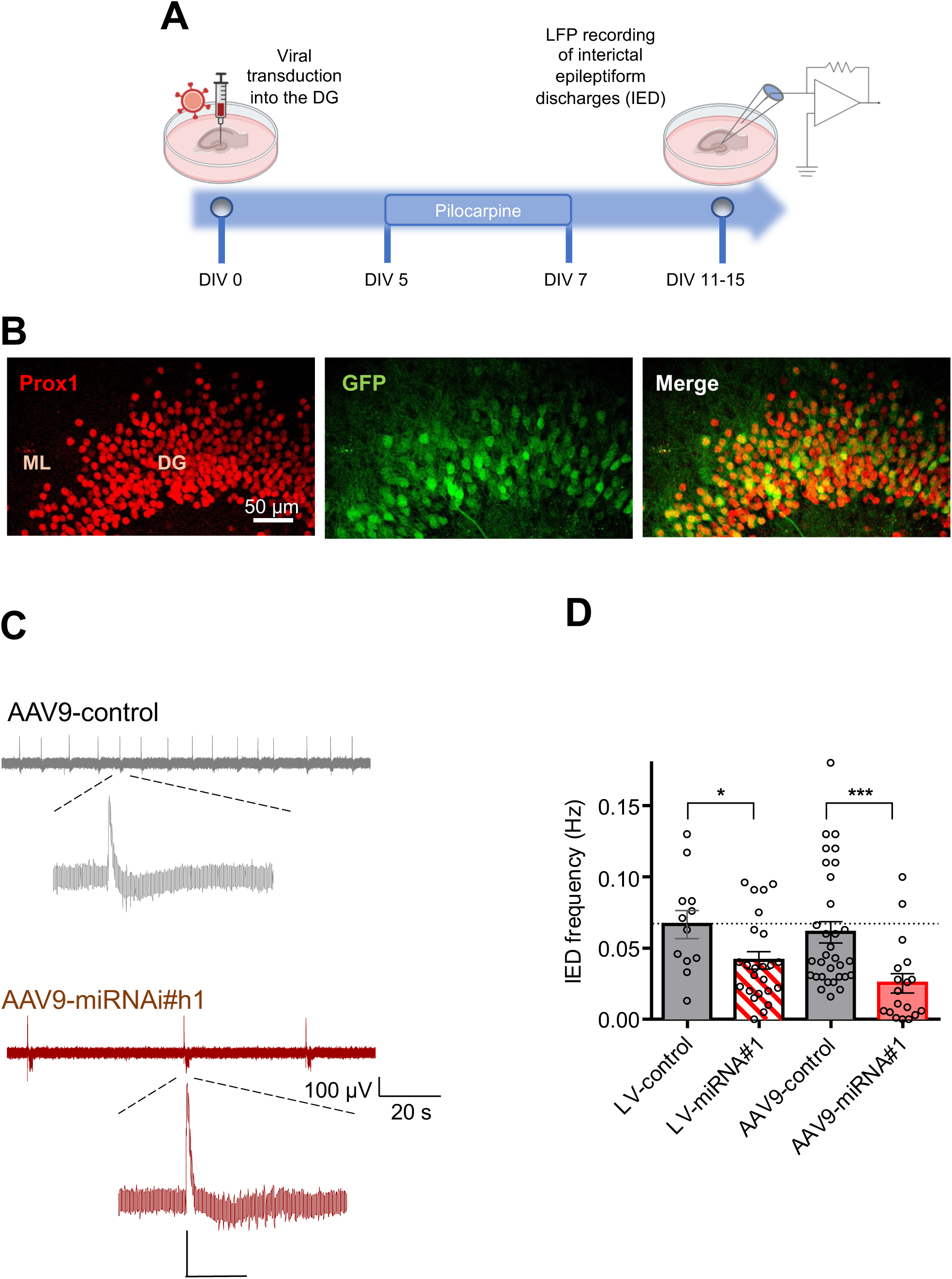
GluK2 RNA silencing suppresses epileptiform activities in mouse organotypic hippocampal slices. **(A)** Illustration of the experimental design and timeline for mouse organotypic slices treated with pilocarpine, transduced with viral vectors and LFP recordings **(B)** Photomicrographs showing an example of dentate granule cells (DGCs) transduced by AAV9–miRScramble (AAV9-control) in a mouse hippocampal organotypic slice using double immunostaining with anti-Prox1 (left), and anti-GFP (middle) antibodies; merged images (right); note that 30% DGCs (prox1 positive cells) are transduced with AAV9-control (GFP positive neurons); molecular layer (ML), scale bar: 50 µm. **(C)** Representative traces of LFPs recorded in mouse hippocampal organotypic slices transduced with either AAV9-control (top left) or AAV9-miRNA#1 (bottom left); enlarged IEDs are shown in the right panel. **(D)** Graph representing the frequency of IEDs in slices transduced with various viral constructs tested: LV-miRNA#1 (n=24 slices from 6 mice) vs LV-control (n=12 slices from 3 mice), *, p=0.0246 by Mann Whitney U test; AAV9-miRNA#1 (n=18 slices from 4 mice) vs AAV9-control (n=32 slices from 8 mice), ***, p=0.0001 by Mann Whitney U test.

**Figure 4:**
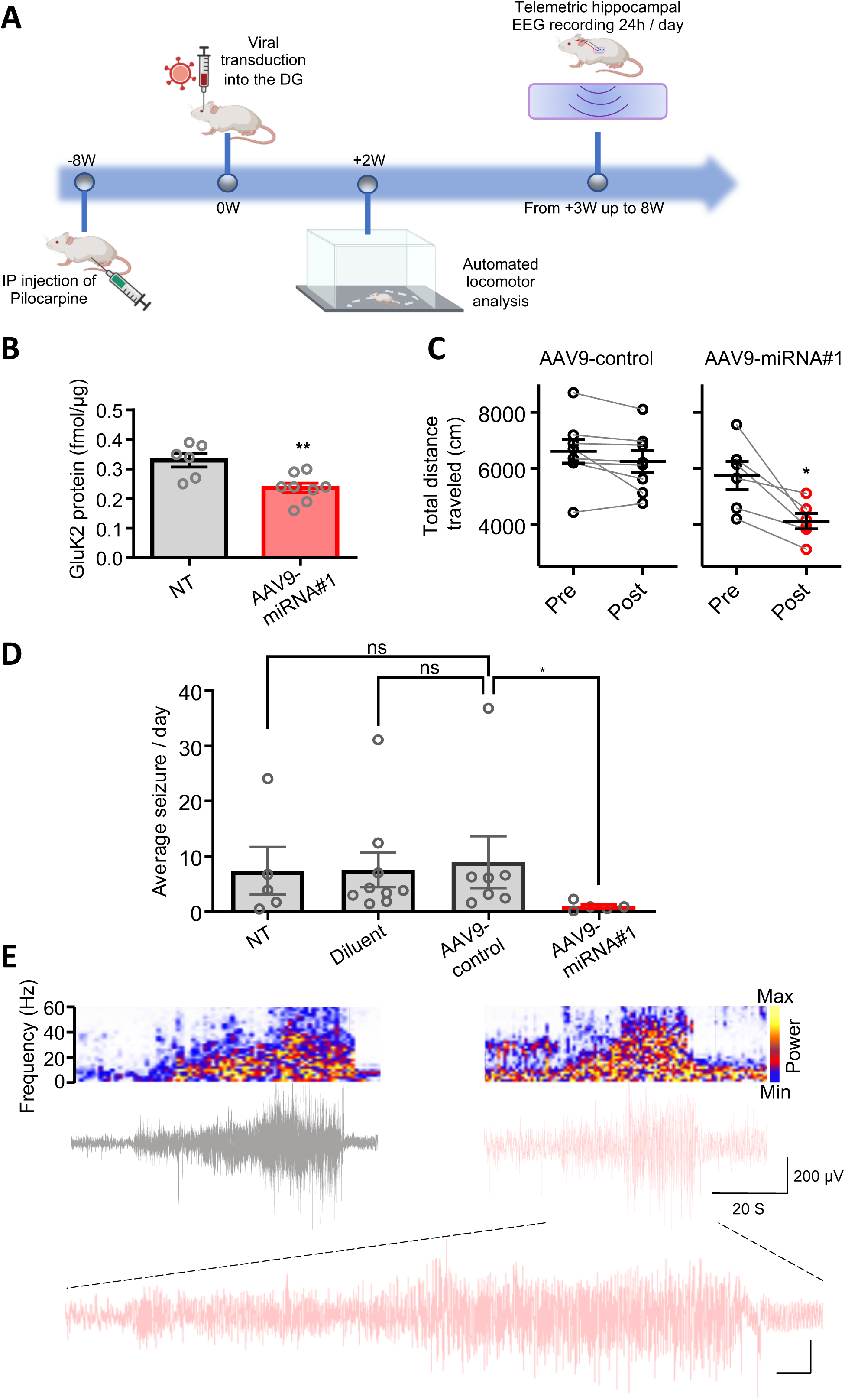
GluK2 RNA silencing suppresses epileptiform activity in a mouse model of TLE. **(A)** Illustration of the experimental design and timeline for a pilocarpine mouse model of TLE, transduced with viral vectors in the hippocampus, tested for locomotion in an open field and recorded with EEG in the hippocampus. **(B)** Absolute quantification of GluK2 by LC-PRM proteomics in brain samples from epileptic pilocarpine mice either non-treated (n 6 hippocampi, 3 mice) or injected with the AAV9-miRNA#1 vector in the hippocampus (n=8 hippocampi, 4 mice), **, p 0.0054, Unpaired t-test. **(C)** Graphs representing the total distance travelled by epileptic mice in the open field before and after treatment with either AAV9-control (p=0.0781, Wilcoxon matched-pairs signed rank test) or AAV9-miRNA#1 (*, p=0.0313, Wilcoxon matched-pairs signed rank test). **(D)** Graph representing the average number of seizures per day in non-treated (NT) chronic epileptic mice (n=5), or chronic epileptic mice treated with either diluent (n=9) or AAV9-control (n=7) or AAV9-miRNA#1 (n=5),*, p<0.05 by One-way Kruskal-Wallis test). **(E)** Examples of seizures recorded in the dentate gyrus of epileptic mice injected with either AAV9-control (grey, left) or AAV9-miRNA#1 (pink, right) and their respective time-frequency spectrograms (above).

**Figure 5:**
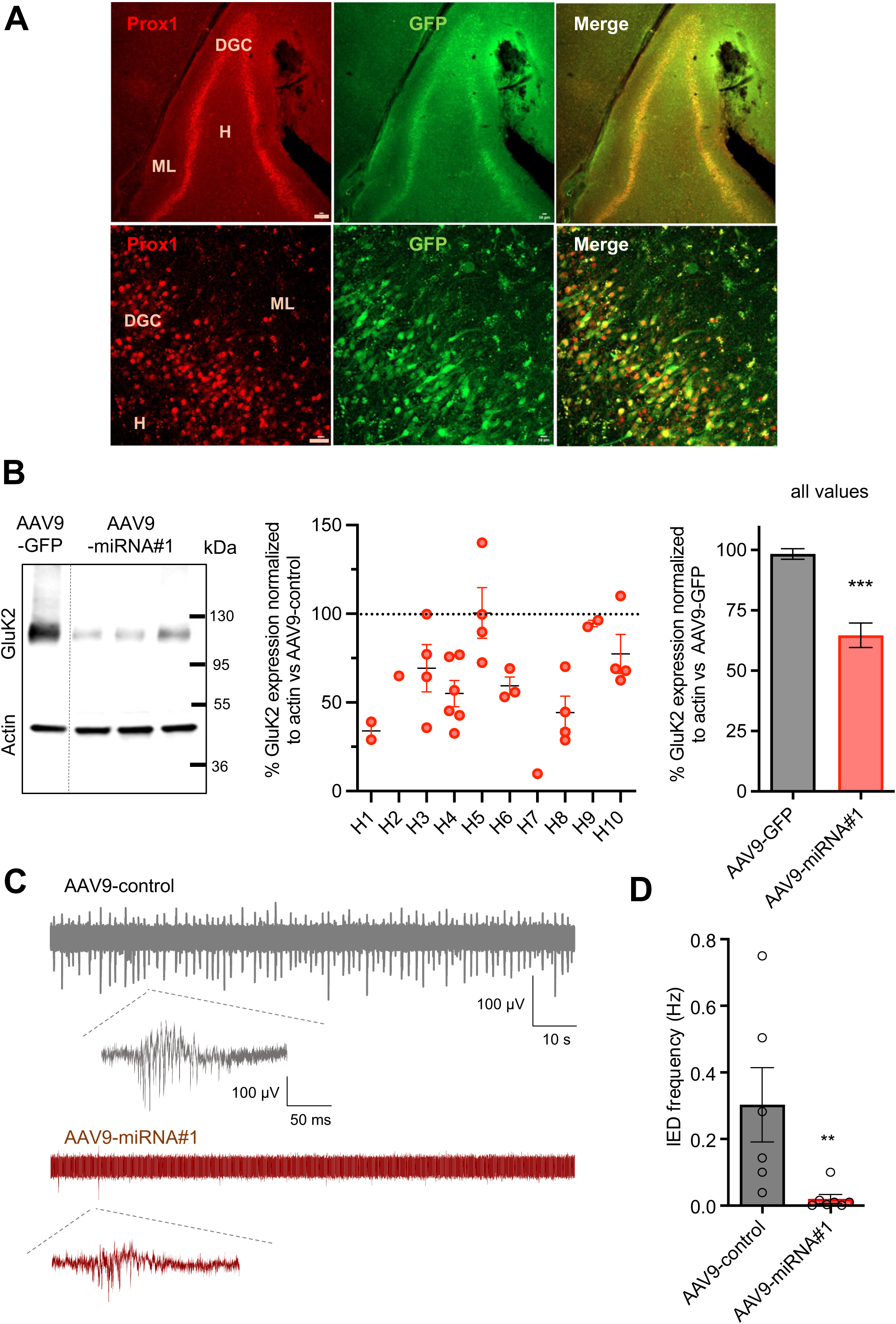
GluK2 RNA silencing suppresses epileptiform activity in human epileptic hippocampus. **(A)** Photomicrographs showing a representative example of DGCs transduced with AAV9-miRScramble (AAV9-control) in hippocampal organotypic slices from a patient with TLE, using double immunostaining with anti-Prox1 (left), and anti-GFP (middle) antibodies. Merged images (right) indicate that about half of DGCs (Prox1 positive cells) were transduced with AAV9-control (GFP positive cells); molecular layer (ML), hilus (H), scale bars: top, 150 µm and bottom, 30 µm. **(B)** Western blot analysis of human organotypic slices transduced with AAV9-miRNA#1 in reference to slices transduced with AAV9-control from the same patients and normalized to β actin; the left panel shows the western blot; middle panel shows individual slices from 10 patients, and the right panel shows the mean values (+/-SEM also shown in table 2) obtained with all slices transduced with either AAV9-miRNA#1 or AAV9-control. (***, p<0.0001 by Mann-Whitney U test). **(C)** Representative traces of LFPs recorded in hippocampal organotypic slices derived from an epileptic patient and transduced with either control AAV9-control (top) or AAV9-miRNA#1 (bottom); enlarged IEDs are shown below. **(D)** Graph representing the frequency of IEDs in organotypic slices from epileptic patients (4 patients) treated with either AAV9-control (n=6 slices) or AAV9-miRNA#1 (n=7 slices)(**, p=0.0029 by Mann-Whitney U test).

**Table 1.** detail and titer of viral vectors used in the study (genome copies GC/ml)

Tables and their legends
| Virus name | description | concentration |
| --- | --- | --- |
| **LV-control** |  |  |
| LV-GFP | LV.hSYN1.GFP | 8.0E+7 GC/ml |
| LV-shScramble | LV.hSYN1.TdTomato.U6.shScramble | 4.6E+7 GC/ml |
| **AAV9-control** |  |  |
| AAV9-GFP | AAV9.hSYN1.GFP | 4.4E+14 GC/ml |
| AAV9-miRScramble | AAV9.hSYN1.GFP.miR30.Scrumble | 9.0E+12 GC/ml |
| AAV9-CAG.EGFP | AAV9.CAG.GFP | 2.0E+13 GC/ml |
| **Test items** |  |  |
| LV-RNA#1 | LV.hSYN1.TdTomato.U6.shRNA#1 | 7.9E+7 GC/ml |
| LV-RNA#2 | LV.hSYN1.TdTomato.U6.shRNA#2 | 8.4E+7 GC/ml |
| LV-RNA#3 | LV.hSYN1.TdTomato.U6.shRNA#3 | 3.3E+8 GC/ml |
| LV-miRNA#1 | LV.hSYN1.TdTomato.miR30.RNA#1 | 3.0E+8 GC/ml |
| **AAV9-miRNA#1** | **AAV9.hSYN1.miR30.RNA#1** | **9.0E+12 GC/ml** |

**Table 2.** Human patients with drug-resistant TLE and subjected to surgical hippocampal resections, used in the study for western blot analysis.

| # patient | Sex | Age | Type (consensus ILAE 2011) | GluK2 decrease vs Control (mean) | SEM |
| --- | --- | --- | --- | --- | --- |
| 1 | M | 56 | Ic | 66% | 5% |
| 2 | F | 24 | unidentifiable | 35% | - |
| 3 | F | 13 | IIa | 31% | 13% |
| 4 | M | 47 | IIIa | 45% | 7% |
| 5 | F | 60 | IIIa | 0% | 14% |
| 6 | M | 11 | Ic | 41% | 5% |
| 7 | M | 18 | IIa | 90% | - |
| 8 | F | 51 | IIb | 56% | 9% |
| 9 | F | 37 | IIa | 5% | 2% |
| 10 | M | 31 | IIIc | 23% | 11% |

### Neuronal cell cultures

Dissociated hippocampal neurons from E18 Sprague–Dawley rat embryos or cortical neurons were prepared from P0-1 C57Bl6J mice (Janvier Labs, France) and GluK2 ko mice^25^. 550,000 neurons were seeded in 6 well plates coated with 1 mg/mL polyLysine were cultured in conditioned Neurobasal medium supplemented with 2 mM L-glutamine and NeuroCult SM1 Neuronal supplement (STEMCELL Technologies) half renewed every 3 days with 3.4mM AraC. After 2-3 days *in vitro* (DIV2-3), vector preparations were added at Multlipicity of Infection (MOI) values of 3 for LV constructs and 7.5E+04 for AAV9 constructs. Ten days after transduction, neuronal cultures (DIV12-13) were scraped into lysis buffer (50 mM HEPES, 100 mM NaCl, 1% Glycerol, 0.5% n-Dodecyl β-D-maltoside, pH 7.2; anti-protease (Calbiochem) and anti-phosphatase (Pierce) cocktails) for western blotting analysis.

### Mouse hippocampal organotypic slices

Organotypic slices (also termed “mouse slice cultures”) were prepared from WT Swiss mice as previously described^20^. At DIV0, 1 µl of phosphate-buffered saline containing the LV or AAV9 viral vectors (LV-shScramble, LV-miRNA#1, AAV9-miRNA#1, and AAV9-control (either AAV9-miRScramble or AAV9-GFP) was applied directly to slices (see Table 1 for titers). Pilocarpine (0.5 µM) was added to the medium at DIV5 and removed at DIV7. All organotypic slices that were transduced and survived the culture process were included in the study without any prior selection. Electrophysiological recordings of Interictal-like Epileptiform Discharges (IEDs) were performed between DIV11 and DIV15.

### Acute and cultured hippocampal slices from TLE patients

Acute and organotypic slices (also termed “human slice cultures”) (thickness 350 μm) were prepared from surgical hippocampal resections from 16 patients (11 – 60 years old) diagnosed with drug-resistant TLE (AP-HM, Hôpital de La Timone, Marseille, France; CHU Pellegrin, Bordeaux, France) as previously described^26, 27^. At DIV 1, slices were transduced with either AAV9-control vectors (AAV9-miRScramble or AAV9-GFP) or AAV9-miRNA#1 applied directly to slices (1.8-4.5E+10 per slice). Electrophysiological recordings were performed either on the same day (acute slices) or between DIV10 and 14. Prior to recordings, acute slices were transferred in oxygenated normal cerebrospinal fluid (ACSF) at room temperature (>1 hour). The ACSF contained (in mM, Sigma-Aldrich): 126 NaCl, 3.5 KCl, 1.2 NaH_2_PO_4_, 26 NaHCO_3_, 1.3 MgCl_2_, 2.0 CaCl_2_, and 10 glucose (pH 7.4). For western blotting analysis, organotypic slices were snap-frozen in liquid nitrogen and lysed in RIPA buffer (Sigma) added with a protease (Calbiochem) and phosphatase (Pierce) inhibitor cocktails with 3 beads using a high-speed benchtop homogenizer FAST PREP24 (MP Biomedicals).

### Electrophysiological recordings and analysis

Mouse and human acute and organotypic slices were individually transferred to a recording chamber maintained at 30-32°C and continuously perfused (2-3 ml/min) with oxygenated (95% O_2_ and 5% CO2) ACSF (physiological conditions), or ACSF containing 5 µM gabazine, or ACSF containing 5 µM gabazine and 50 µM 4-AP. In mouse hippocampal slices, spontaneous IEDs were examined using local field potential recordings in the presence of 5 µM gabazine as previously described^20^. In human hippocampal slices, the integrity of the epileptic microcircuit to generate spontaneous IEDs was systematically probed under hyperexcitable conditions (5 µM gabazine and 50 µM 4-AP); spontaneous IEDs examined by local field potential (LFP) recordings consisted of events including multi-units and population spikes (30 – 2200 ms duration) as previously described^27, 28^ (see also **Fig 1** and **Fig 5**). For experiments with UBP310, slices were continuously monitored in the presence of gabazine and 4-AP. For experiments with viral vectors, organotypic slices which responded positively with ongoing spontaneous IEDs in the presence of gabazine and 4-AP, were then switched back to standard ACSF and washed at least for 25 min before analysis in these physiological conditions. Signals were analyzed off-line in a blinded manner using Clampfit 9.2 (PClamp) and MiniAnalysis 6.0.1 (Synaptosoft, Decatur, GA).

### Western blotting

The WB experiments were performed in a blinded manner with respect to the treatment with viral vectors. Ten µg of total proteins quantified by Pierce^TM^ BCA protein assay kit (ThermoScientific) were loaded on 4-15% gradient SDS–PAGE gels and transferred to nitrocellulose membranes (Trans-Blot turbo, Bio-Rad). Immunoblotting of rodent cultures was detected by chemiluminescence with clarity Western ECL on ChemiDoc Touch system with Image Lab software (Bio-Rad) using a rabbit anti-GluK2/GluK3 (Merck-Millipore 04-921; 1:2000), a mouse anti-β actin (Sigma A5316; 1:5000) and appropriate HRP-conjugated secondary antibodies (Jackson ImmunoResearch; 1:5000). Immunoblotting of human organotypic slices was detected in fluorescent on Li-COR with Image studio software (Biosciences GmbH) using a rabbit anti-GluK2 (Abcam ab124702; 1:2000), a mouse anti-ý actin (Sigma A5316; 1:5000) in Odyssey blocking buffer TBS (Li-COR) and appropriate 800nm fluorophore-conjugated secondary antibodies (IRDye 800; 1:15000). The viability of slices was confirmed by total proteins stained with Revert 700nm kit (Li-COR). The intensity of the signal for GluK2 or gluK2/GluK3 of each lane was normalized to the β actin signal and thereafter to the control condition.

### Real-time RT-qPCR

The qPCR experiments were performed in a blinded manner with respect to the treatment with viral vectors. Total RNA was extracted from DIV12 infected mouse cortical neurons using the Direct-zol RNA microprep (Zymo). Ten µg were reverse-transcribed into cDNA by Maxima First Strand cDNA Synthesis Kit (Thermofischer). Quantitative real-time qPCR was performed in a LightCycler480 real-time PCR system (Roche) using TaqMan Gene Expression Assays (Thermo Fisher Scientific) for mouse *grik1* (Mm00446882_m1)*, grik2* (Mm01181234-m1)*, grik3 (Mm01179722_m1), grik4 (Mm00615472_m1), grik5 (Mm00433774_m1)*, *gria1* (Mm00433753_m1), *gria2* (Mm01220174_m1), *grin2a* (Mm00433802_ m1), *grabrb2* (Mm00549788_s1), *gabrb3* (Mm00433473_m1) and *gapdh* (Mm99999915-g1) as a housekeeping gene^29, 30^.

### Immunostaining of mouse and human organotypic slices

Slices fixed in PFA 4% after recording were permeabilized in blocking solution containing 5% normal goat serum (NGS) (mouse slices) or 10% horse serum (HS) (human slices) and PBST (PBS + 0.5% Triton) for 2 hours at room temperature, and were incubated with anti-GFP antibody (Abcam, AB-300798, 1:1000) and either with anti-Prox1 antibody (Millipore ab5475, 1:2000 (mouse slices:) or R&D system AF2727, 1:100 (human slices)) for 48 hours at 4°C. Slices were then incubated for 2 hours with Alexa Fluor conjugated secondary antibodies (Invitrogen 1:500) and coverslipped in Fluoromount (Thermo Fisher). Fluorescence images were acquired using a LSM800 confocal microscope (Zeiss) using 10X/0.3NA and 20X/0.8NA objectives. Images were processed using NIH ImageJ software.

### AAV9 stereotaxic injection in non-epileptic and chronic epileptic mice

Chronic epileptic mice (Swiss, 30-40 g, purchased from Janvier and Charles River Labs) were generated using an adapted pilocarpine-induced status epilepticus (SE) model^31^. Control (non-epileptic) and epileptic mice (>2 months after SE) were bilaterally injected with AAV9 constructs into the dorsal and ventral hippocampus (AP -1.8 mm, ML ±1 mm, DV – 2 mm, and AP -3.3 mm, ML ± 2.3 mm, DV – 2.5 mm) as previously described_32_. A volume of 9 E+9 GC in 1 µl/injection site x 4 injection sites of either AAV9-control (AAV9-miRScramble or AAV9-GFP) or AAV9-miRNA#1 constructs was injected with a Hamilton syringe (rate: 0.2 µl/minute). In order to evaluate the spread of transduction along the hippocampus, immunohistochemical detection of GFP following stereotaxic injections of AAV9-hSyn1-GFP was performed in non-epileptic mice. After 3 weeks, mice were deeply anesthetized with ketamine/xylazine and transcardially perfused with normal saline (0.9% NaCl). Brains were dissected and post-fixed in 4% paraformaldehyde for 48 hours followed by cryoprotection in 30% sucrose solution. Cryoprotected brain hemispheres were sectioned in the sagittal plane at 50 µm on OCT compound (Tissue Tek, Torrance3, CA, USA) using a Leica freezing microtome. Sections were then incubated with primary antibodies – chicken anti-GFP (#NB100-1614, Novus Biologicals, Centennial, CO, USA; 1:1000); mouse anti-NeuN (#ab104224 Abcam, Cambridge, MA, USA; 1:1000). Confocal images were acquired using a Zeiss LSM 900 Airyscan 2. A 63X objective was used to acquire tiled images of CA1, CA2, CA3, and dentate gyrus/hilus in a single z-plane, and all analysis was conducted using the HALO Image Analysis Software (Indica Labs, Albuquerque, NM, USA). An artificial intelligence based nuclear-segmentation classifier was trained via manual segmentation to identify individual cells based on NeuN. ROIs were drawn around each region (CA1, CA2, CA3, dentate gyrus granular cells, and hilus) based on cytoarchitectural characteristics in relation to demarcations in a mouse brain atlas (Paxinos and Franklin, 2007). Using the Highplex FL module, GFP intensity thresholds were set for each ROI and colocalization of GFP with classified NeuN+ cells was used to calculate the percent of neurons that were positive for GFP. The average GFP intensity of all classified neurons was also determined (e.g. the GFP luminance of each NeuN+ cell, including those that were categorized as both GFP-positive and GFP-negative). The % of GFP-positive neurons in reference to NeuN-positive neurons was measured in 4 mice close to the injection site using this procedure was: 76.8 ± 5.6% in the DG, 87.5 ± 8.4% in the hilus, 92.2 ±7.8% in CA1, 88.5 ± 6.1% in CA2, and 96.4 ± 2.2 in CA3. Therefore, all areas of the hippocampus are strongly transduced with AAV9 in the vicinity of the injection site.

### Quantification of GluK2 in mouse brain samples using mass spectrometry

In order to quantify GluK2 protein from brain samples with high accuracy, we used liquid chromatography mass spectrometry and a parallel reaction monitoring (LC-PRM) workflow^33^. Brain samples from pilocarpine-treated mice were homogenized and denatured (Precellys Evolution Homogenizer, Bertin Instruments) then reduced and alkylated using Biognosys’ Reduction and Alkylation solution. Samples were digested overnight with sequencing grade trypsin (Promega) at a protein:protease ratio of 50:1. Peptides were desalted using 96-well µHLB plate (Waters) and dried down using a SpeedVac system. Peptides were resuspended in 1 % acetonitrile and 0.1 % formic acid and spiked with Biognosys’ iRT kit calibration peptides. Peptide concentrations in mass spectrometry ready samples were measured using the mBCA assay (Thermo Scientific™ Pierce™). Three stable isotope labeled reference peptides were spiked into the final peptide samples at known concentrations (Vivitide, the quality grade of the reference peptides was ±10% quantification precision, >95% purity; purity of peptide TVTVVYDDSTGLIR was 93.4 %). Three peptides representing GluK2 protein and the corresponding SIS reference peptides were monitored for absolute quantification; 3 peptides from mouse proteins were included as a control for sample preparation and loading: GAPDH (Glyceraldehyde-3-phosphate dehydrogenase), PARK7 (Parkinson disease protein 7 homolog), ACTG1 (Actin, cytoplasmic 2). For LC-PRM measurements, 1 µg of peptides per sample was injected to an in-house packed C18 column (PicoFrit emitter with 75 µm inner diameter, 60 cm length, and 10 µm tip from New Objective, packed with 1.7 µm Charged Surface Hybrid C18 particles from Waters) on a Thermo Scientific Easy nLC 1200 nano-liquid chromatography system connected to a Thermo ScientificQ Exactive HF-X mass spectrometer equipped with a standard nano-electrospray source. LC solvents were A: 1 % acetonitrile in water with 0.1 % FA; B: 20 % water in acetonitrile with 0.1 % FA. The LC gradient was 0 – 59 % solvent B in 54 min followed by 59 -90 % B in 12 sec, 90 % B for 8 min (total gradient length was 67 min). A scheduled run in PRM mode was performed before data acquisition for retention time calibration^34^. The data acquisition window per peptide was 6.7 minutes. Signal processing and data analysis were carried out using SpectroDive™ 11.6(Biognosys) for multiplexed MRM/PRM data analysis based on mProphet^35^. A Q-value filter of 1% was applied.

### Locomotor activity in non-epileptic and chronic epileptic mice

Locomotion was assessed in chronic epileptic mice (>2 months after SE) 1 week before and 2 weeks after AAV9 injection. Mice were housed at room temperature (20–22◦C) in a 9:00 – 18:00 light/ dark cycle with ad libitum access to food and water. Mice were transferred to the test room 1 day prior to experimentation for habituation. All materials which have been in contact with the animal during testing were washed thereafter in order to prevent olfactory cues. Spontaneous exploration behavior was tested with the open field test^36^. The mice were placed into the center of a 50 × 50 × 50 cm blue polyvinyl chloride box, and the trajectories were recorded with a video camera connected to a tracking software EthoVision Color (Noldus, Netherlands) to automatically measure the total distance travelled by the mice during 10 min. Age-matched non-epileptic naïve mice were also evaluated for locomotion.

### EEG recordings in chronic epileptic mice

Chronic epileptic mice (>2 months after SE) were implanted with one depth wire DSI electrode as described previously^20^, 3 weeks after AAV9 injection. The recording electrodes were placed stereotaxically into the DG (coordinates from bregma: AP -2.55 mm, ML +1.65 mm, DV -2.25 mm). EEG was analysed in a blinded manner (using the Ponemah software DSI, St. Paul, MN). Seizures were defined as rhythmic (> 4 up to 60 Hz, see Fig 4D) and prolonged spike trains with an amplitude of at least twice the EEG baseline and a duration of at least 8 second^37, 38^; the time-frequency spectrogram of EEG was performed using a Fast Fourier Transform (FFT) algorithm with a sliding 1s-Hanning windows (‘periodogram’ function of DSI).

### Statistical analysis

All values are given as means ± SEM unless otherwise specified. Statistical analyses were performed using Graphpad Prism 7 (GraphPad Software, La Jolla, CA). The Shapiro–Wilk test was used to determine normality of the data. The parametric Student’s t test (paired and unpaired, two-tail) was used to compare normally distributed groups of data. The Mann-Whitney test (unpaired data, two-tail) and Wilcoxon Signed Rank test (paired data) were used for non-normally distributed data. For the comparison of multiple groups, one or two-way ANOVA test were used. P-values and statistical tests are given in the figure legends, except for those not derived from figures. The level of significance was set at p<0.05. Group measures are expressed as mean ± SEM; error bars also indicate SEM. All *in vitro* and *in vivo* experiments were performed blind to the identity of the viral vector used.

## Results

### Pharmacological inhibition of KARs reduces interictal-like epileptiform activity in hippocampal slices from patients with TLE

To determine the role of KARs in epileptiform network activity, a proof-of-concept study was conducted using the KAR antagonist UBP310^20, 39^ in the DG of drug-resistant TLE patient-derived hippocampal slices. Local field potentials (LFPs) recorded in organotypic hippocampal slices between DIV 10 and DIV21 in the presence of 5 µM gabazine and 50 µM 4-AP demonstrated spontaneous recurrent Interictal-like Epileptiform Discharges (IEDs) (**Fig 1B**, top traces). UBP310 at a concentration (5 µM) which selectively targets KARs over other ionotropic glutamate receptors (iGluRs), significantly diminished the frequency of IEDs by approximately 50% (from 0.057 ± 0.010 Hz at baseline to 0.027 ± 0.008 Hz in the presence of UBP310, n=9 slices, 4 patients, p<0.01, **Fig 1B-D**) without a significant change in their duration (1.29 ± 0.19s before versus 1.57 ± 0.35s after UBP310, p=0.0625, Wilcoxon matched-pairs signed rank test, n=9 slices, 4 patients). It is worth noting that the frequency of IEDs remained stable over the duration of the recording in the absence of UBP310 (**Fig 1C**). Similar experiments were conducted in acute hippocampal slices from TLE patients; UBP310 (5 µM) reduced the frequency of IEDs by around 50% (from 0.045 ± 0.011 s at baseline to 0.027 ± 0.009 Hz in the presence of UBP310, p=0.0020, Wilcoxon matched-pairs signed rank test, n=10 slices), without a significant change of their duration (0.43 ± 0.09 s before versus 0.37 ± 0.08s after UBP310, p=0.3203, Wilcoxon matched-pairs signed rank test, n=10 slices). In summary, pharmacological inhibition of KARs reduces IEDs in hippocampal slices from patients with TLE.

### Design and validation of viral vectors for the downregulation of GluK2 by RNA interference

Because UBP310 lacks subunit selectivity^40, 41^, we rationally developed a virally-delivered RNA interference (RNAi) approach to selectively downregulate GluK2.Three RNAi sequences (RNA#1, #2, and #3) were designed against the human *grik2* gene and evaluated by generating LV vectors expressing shRNAs and by transducing cultured rat hippocampal neurons. The transduction rate of LV vectors was 84 ± 4% in rat hippocampal neuronal cultures, as estimated by the quantification of GFP-expressing neurons. Transduction of neurons with LV vectors expressing shRNA#1 was the most efficient in reducing total GluK2/GluK3 protein levels **(Fig 2A** and **2B)** (shRNA#1: decrease by 30 ± 7%, n=11, p<0.0001, shRNA#2: decrease by 22 ± 7%, n=4, p>0.05, shRNA#3: decrease by 17 ± 10%, n=3, p>0.05 vs. LV-GFP n=13) when compared with LV-GFP control constructs; hence shRNA#1 was investigated further. Viral vectors were further designed and tested for the expression of the engineered RNAi corresponding to shRNA#1 (**Fig 2C**) using a human miR30 structure (miRNA#1) under the control of hSYN1 promoter, either in the context of LV or AAV9 vectors (**Fig 2B**). Transduction of rat hippocampal cultures with these viral constructs expressing miRNA#1 significantly reduced GluK2/GluK3 protein levels by 36 ± 7% (n=12, p< 0.0001) with LV-miRNA#1 and by 35 ± 5% (n=15, p< 0.0001) with the AAV9-miRNA#1 as compared with LV-GFP and AAV9-GFP respectively (**Fig 2B**). As expected, the LV-shScramble (109± 11%, n=4, p>0.05) and AAV9-miRScramble (114 ± 5%, n=5, p>0.05) showed no significant reduction when compared with LV-GFP (102 ± 4%, n=13) and AAV9-GFP (101 ± 1%, n=16) vectors, respectively (**Fig 2B**). The RNAi sequences designed using the human *grik2* mRNA sequence were complementary to rat *grik2* mRNA, however they had one mismatch with the mouse *grik2* mRNA sequence (**Fig 2C**). Before further functional analysis *ex vivo* and *in vivo* in mice, we validated the efficiency of AAV9-miRNA#1 to downregulate GluK2 in mouse cortical neurons *in vitro*. Ten days after transduction of mouse cortical neurons with AAV9-GFP, 89 ± 2% of neurons expressed GFP (mean of n=6 independent neuronal cultures) (**Fig 2D**). In the absence of a suitable antibody strictly selective for the GluK2 subunit in rodents, we used again the GluK2/3 antibody; by performing western blot analysis of cortical cultures from GluK2 ko mice, we could evaluate that GluK2 accounts for more than 75% of the total labeling intensity (**Fig 2E**). Transduction of mouse cortical neurons with AAV9-miRNA#1 decreased GluK2/GluK3 protein levels by 37 ± 3% (n=23, vs. AAV9-control, 101 ± 2%, n=21, p<0.0001) (**Fig 2E**). The decrease in GluK2/GluK3 protein level was associated with a decrease of *grik2* mRNA measured by qPCR by 30 ± 2 % (vs AAV9-control, 99 ± 5%, n=9, p<0.05) (**Fig 2F**). We tested by qPCR the specificity of the downregulation of *grik2* mRNA by comparing in the same samples the effect of AAV9-miRNA#1 on the other KAR subunit mRNAs (*grik1*, *grik3*, *grik4* and *grik5* mRNA), as well as on major AMPA, NMDA and GABA_A_ receptor subunits (*gria1*, *gria2*, *grin2a*, *gabr2*, *gabr3*) **(Fig 2F).** No significant difference was found for any of these mRNA (n = 9 from 3 independent neuronal cultures, p>0.1) (**Fig 2F**). These data clearly highlight the selectivity of AAV9-miRNA#1 treatment for *grik2* mRNA and GluK2 KAR subunit.

### GluK2 silencing suppresses interictal-like epileptiform activities in mouse organotypic hippocampal slices and chronic seizures in a mouse model of TLE

The ability of LV or AAV9-miRNA#1 to reduce the frequency of IEDs was evaluated in organotypic mouse hippocampal slices exposed to pilocarpine for 2 days; these slices displayed recurrent mossy fibers and KAR-mediated IEDs^20, 42^, as shown in animal models of TLE^21^. These pilocarpine-treated organotypic slices were successfully transduced with AAV and LV vectors, and in particular DGCs, as indicated by the colocalization of GFP+ DGCs and Prox1 staining following transduction with AAV9-GFP vectors (**Fig 3B**). To ensure the recording of reliable and stereotyped spontaneous IEDs in this *in vitro* model, LFP recordings were performed in the continuous presence of a GABA_A_-R antagonist (5 µM gabazine) as previously described^20, 43^. In this condition, we showed that transduction of mouse organotypic slices with LV-miRNA#1 (3E+5 GC/slice) strongly reduced the frequency of IEDs measured at DIV11-DIV15 when compared with controls (LV-shScramble) (0.041 ± 0.006 Hz with LV-miRNA#1, n=24; 0.066 ± 0.009 Hz with LV-shScramble, n=12, p<0.05) (**Fig 3C, D**). A similar reduction of the frequency of IEDs (when compared with control conditions) was observed with AAV9-miRNA#1 (9E+9 GC/slice) (0.025 ± 0.007 Hz with AAV9-miRNA#1, n=18; 0.061 ± 0.007 Hz with AAV9-control, n=32, p<0.0001) (**Fig 3C, D**).

The efficacy of AAV9-miRNA#1 in reducing epileptic activity *in vivo* was determined using the pilocarpine mouse model of TLE. AAV9-miRNA#1 and AAV9-control vectors were delivered to the hippocampus of chronic epileptic mice (**Fig 4A**). A high rate of neuronal transduction was observed in mouse hippocampal neurons when AAV9-GFP was injected *in vivo* in the hippocampus of control non-epileptic mice (see methods). We evaluated the impact of AAV9-miRNA#1 *in vivo* delivery on the level of GluK2-containing KARs by using a proteomic approach (LC-PRM), 7 weeks after viral injection, in a whole hippocampus (hence including regions at various distances from the injection sites). This allowed obtaining an absolute quantification of GluK2 protein in homogenates of hippocampal samples from epileptic mice, either treated with AAV9-miRNA#1 or non-treated (0.24 ± 0.016 fmol/µg with AAV9-miRNA#1, n=8 hippocampus, 4 mice; 0.33 ± 0.023 fmol/µg in non-treated condition n=6 hippocampus, 3 mice, p=0.0054, Unpaired t-test) (**Fig 4B**), which corresponds to a knock-down of GluK2 by around 30% in the whole hippocampus.

We next assessed whether knock-down of GluK2 in epileptic mice was able to improve hyperlocomotion (**Fig 4A, C**), a well-described behavioral alteration associated with epilepsy^36^. Chronic epileptic mice displayed pathological hyperlocomotor activity when compared to non-epileptic naïve mice (total distance traveled in 10 min: 6583 ± 421 cm in 8 chronic epileptic mice versus 2945 ± 114 cm in 8 naïve mice, p <0.001, Mann Whitney test). The treatment with AAV9-miRNA#1, injected bilaterally in the DG of chronic epileptic mice (3.6E+10 GC/brain), reduced locomotor activity towards values in naïve mice (from 5746 ± 501 cm before injection to 4118 ± 278 cm 15 days after injection, n=6, p<0.05) **(Fig 4C).** No significant reduction of hyperlocomotor activity was observed after treatment of chronic epileptic mice with AAV9-control (either AAV9-miRScramble or AAV9-GFP constructs, 3.6E+10 GC/brain) (from 6583 ± 421 cm to 6218 ± 383 cm, 15 days after injection, n=8, p>0.05) (**Fig 4C**). Three weeks after viral injection, the same group of mice were then monitored with telemetric EEG recordings in the DG (**Fig 4A, D**). Treatment with AAV9-miRNA#1 significantly reduced the number of chronic seizures (2.55 ± 1.45 seizures/day with AAV9-miRNA#1, n=6; 16.55 ± 5.2 seizures/day with AAV9-control constructs, n=8, p<0.05) as well as the cumulative duration of the seizures recorded over 5 days (6.9 ± 3.2 min with AAV9-miRNAi#1, n=6; 45.4 ± 12.9 min with AAV9-control constructs, n=8, p<0.05) without any significant change neither in their duration (39.4 ± 9.4 s with AAV9-miRNAi#1, n=3; 34.9 ± 4.2 s with AAV9-control constructs, n=3, p=0.89, Mann Whitney test) nor in their time-frequency spectrogram (**Fig 4E**). To confirm the robustness of the treatment on seizure activity with AAV9-miRNA#1 over time, a subsequent group of mice was followed up beyond one month after treatment (6-7 weeks after viral injection). In this condition, we also observed a significant reduction in the number of seizures in mice recorded 42-49 days after the injection (0.9 ± 0.3 seizures/day with AAV9-miRNA#1, n=5, p<0.05 by One-way Kruskal-Wallis test; 9.0 ± 4.7 seizures/day with AAV9-control constructs, n=7; 7.6 ± 3.2 seizures/day with diluent, n=9; 7.4 ± 4.3 with non-treated condition, n=5)(**Fig 4D**). The sustainability of the AAV9-miRNA#1 treatment was further substantiated by the quantitative proteomic analysis of GluK2 expression, which was also performed 7 weeks post-treatment (see above, **Fig 4B**). Taken together, these data demonstrate that GluK2 knock-down suppresses IEDs in an *in vitro* model as well as seizures in an *in vivo* mouse model of TLE, and it further indicates that this effect is sustained for at least several weeks after a single viral injection in mice.

### GluK2 silencing suppresses interictal-like epileptiform activities in hippocampal slices from patients with TLE

The therapeutic potential of gene therapy targeting GluK2 was assessed using organotypic hippocampal slices from patients with drug-resistant TLE as an *ex vivo* translational disease model. The ability of AAV9 (1.8E+10 GC/slice) to transduce human slice cultures was evaluated with AAV9-GFP vectors. Co-labelling of human organotypic slices with a Prox1 and a GFP antibody revealed that 57.7 ± 3.8 % of DGCs were GFP-positive (n=4 slices from 3 patients) (**Fig 5A**). Slices from 10 patients (**Table 2**) were then transduced with AAV9-miRNA#1 or AAV9-control (1.8-4.5E+10 GC/slice) constructs. Western blotting experiments using a subunit-selective GluK2 antibody suitable for human samples showed that, on average, transducing the slices with AAV9-miRNA#1 significantly decreased GluK2 protein levels by 33 ± 5% with reference to either AAV9-GFP or AAV9-miRScramble constructs (AAV9-control; see methods) (n=31 slices from 10 patients, p<0.0001, Mann-Whitney test) (**Fig 5B**).

The efficacy of AAV9-miRNA#1 in reducing IEDs was assessed in a further set of human organotypic slices. For these, the transduced slices were recorded in ACSF to test the efficacy of AAV9-miRNA#1 under physiological medium conditions. When transduced with an AAV9-control construct (2E+10 GC/slice), these slices displayed frequent spontaneous recurrent IEDs (0.3 ± 0.1 Hz, n=6 slices from 4 patients). In contrast, when slices were treated with AAV9-miRNA#1, we observed a suppression of IEDs (by around 93%; 0.019 ± 0.013 Hz, n=7 slices from 4 patients; p<0.01) (**Fig 5C-D**). These data demonstrate that downregulation of GluK2 by a viral-mediated RNAi strategy is effective in suppressing IEDs in hippocampal tissues from patients with drug-resistant TLE.

## Discussion

Here we describe a novel gene silencing approach for the treatment of drug-resistant TLE, for which novel therapeutic strategies are urgently required. An AAV9-based gene therapy vector was engineered to deliver a miRNA specifically targeting GluK2 containing KARs, which demonstrated the successful inhibition of **IEDs *in vitro* and chronic seizures** *in vivo* in a mouse model of TLE. Most importantly, transduction of the AAV9-anti *grik2* miRNA reduced GluK2 levels and mediated the reduction of spontaneous IEDs in hippocampal organotypic slices from drug-resistant TLE patients.

Pharmacological inhibition of GluK2/GluK5 KARs or genetic removal of GluK2 is efficient in reducing the frequency of epileptic seizures in mouse models of TLE^20^; however, it is important to address whether this approach can be applied to human TLE. In rodents models of TLE, post-synaptic pathological KARs ectopically expressed at recurrent mossy fiber synapses^18, 19, 39, 42^ promote chronic and recurrent seizures^20, 42^. KAR subunits identified in rodents show high homology with KAR subunits expressed in human^44^. Whereas in rodents, both electrophysiological and immunohistochemical experiments demonstrate the presence of ectopically expressed KARs within the inner molecular layer of the DG^18, 42^, this has not been demonstrated in the hippocampus of TLE patients because selective antibodies for either GluK2 or GluK5 are not currently available. However, evidence obtained from high-affinity kainate binding studies implies that GluK2/GluK5, is enriched in the DG inner molecular layer in TLE^14, 44^. Mossy fibers sprouting (a key histopathological feature of the TLE hippocampus)^15^ creates recurrent excitatory synaptic contacts onto DGCs^17^. The mechanisms by which KARs are recruited and stabilized at these aberrant synapses share similarities with the mechanisms operating at mossy fiber CA3 synapses^42^. It is therefore expected that mossy fiber sprouting in human TLE is accompanied by the recruitment of KARs at ectopic pathological synapses. Further work should clarify the subcellular distribution of KAR subunits in the DG of the human TLE hippocampus once appropriate methods and antibodies are available.

Here we have shown that UBP310, an antagonist of KARs, alleviates recurrent IEDs in acute and cultured hippocampal slices from resective epilepsy surgery in TLE patients. This strongly suggests a role for KARs in the generation of epileptic seizures in patients with mesial TLE. UBP310 was initially designed as a GluK1 antagonist^40^ but was subsequently shown to antagonize synaptic KARs comprising GluK2/GluK5 receptors^39^. UBP310 is not a candidate antiepileptic molecule as it lacks subunit selectivity and cannot cross the blood-brain barrier^40, 41^.

Some existing drug therapies for TLE may act via KAR modulation^45–47^. Topiramate is a broad spectrum antagonist of AMPA/kainate receptors^48^, but other potential targets for topiramate have also been described^45^. Perampanel, primarily designed as a non-competitive negative allosteric modulator of AMPA^49^, is also an allosteric modulator of KARs^47^. Topiramate is associated with numerous side effects and was notably included in an FDA alert for increased risk of suicidality^50^. The utility of perampanel is limited by its narrow therapeutic index, and furthermore, it carries a regulatory (boxed) warning for serious psychiatric and behavioral adverse reactions^49, 51^. Finally, because GluK2/GluK5 are highly abundant KARs expressed in most brain regions^52^, a pharmacological approach antagonizing these receptors in all brain regions could have undesired effects on neuronal circuits and associated behaviors^53^.

Instead, we provide strong evidence for an alternative gene silencing strategy that selectively targets GluK2 using miRNA silencing. We identified an RNA silencing sequence that efficiently downregulated GluK2 protein expression in cultured hippocampal neurons both as a shRNA expressed under the control of the U6 promoter or as a miRNA expressed under the control of the neuronal promoter hSYN1. Based on these screening experiments, an AAV9-based vector was engineered to express the most efficient anti GluK2/*grik2* miRNA driven by the neuron-specific hSYN1 promoter. AAV9 was selected as a delivery vector because it efficiently transduces cells in the CNS, including neurons, following intracerebral vector administration^54^. In control experiments, stereotaxic injection of reporter gene expressing control (AAV9-GFP) vectors at two independent sites demonstrated transduction of neurons in the hippocampus without significant off-hippocampal expression of the fluorescent marker. The effect of GluK2 knock-down *in vivo* by AAV9-anti *grik2* miRNA (AAV9-miRNA#1) was investigated in a pilocarpine mouse model of TLE. AAV9-anti *grik2* miRNA was administered by stereotactic bilateral hippocampal injection. The treated epileptic mice were recorded with telemetric EEG over several days displaying recurrent chronic seizures that are drastically reduced in mice transduced with the AAV9-anti *grik2* miRNA (by approximately 85%). This result is in line with the reduced chronic epileptic activity in constitutive GluK2 knockout mice treated with pilocarpine^20^. It further shows that suppressing GluK2 in the hippocampus alone is sufficient to alleviate epileptic activity. Furthermore, it indicates that partial downregulation of GluK2 by approximately 30% on average for the whole hippocampus (as opposed to full knockout of GluK2) remains efficient as an antiepileptic strategy. This is consistent with the notion that KARs are a major contributor to the abnormal detonator features of epileptic DGCs, in particular due to the slow decay kinetics of KAR-EPSCs allowing for summation of depolarization to reach spike discharge threshold^55^. Therefore, lowering the amount of these KARs appears sufficient for the circuit to remain under the threshold for seizure generation.

It is important to underscore the fact that the knockdown of GluK2 *in vivo* is long-lasting, as shown by the LC-PRM proteomic analysis which was performed 7 weeks after *in vivo* viral transduction with AAV9-miRNA#1, and it is in line with the demonstration that the robust effects of the treatment on epileptic activity are also sustained.

Our data in rodents show that GluK2/GluK5 is a potential therapeutic target for TLE; we further tested whether the downregulation of GluK2 using our RNAi approach could efficiently reduce epileptic discharges in the context of the human hippocampus. We levered the possibilities offered by long-term culture of hippocampal slices obtained from surgical resections of patients with drug-resistant TLE. Our data confirmed effective transduction of human slice cultures by AAV9 vectors^27^. Transduction of hippocampal organotypic slices from TLE patients with AAV9-anti *grik2* miRNA resulted in reduced levels of GluK2, and, most importantly in the significant reduction of spontaneous epileptic discharges, in line with the results obtained with UBP310.

Previous studies in mice have established that inhibition of GluK2 synthesis is sufficient to fully abolish GluK2/GluK5 KAR function, since GluK5 by itself does not form functional homomeric channels^56, 57^. Genetic deletion of GluK2 was shown to indirectly downregulate GluK5^56, 57^, probably because GluK5 is not expressed at the cell membrane as a homomeric receptor and is degraded in the absence of the partner subunit GluK2. Hence, the RNAi-based downregulation of GluK2 may suppress the formation of GluK2/GluK5 KARs in the cell membrane. We therefore postulate that in the epileptic hippocampus, RNAi-based downregulation of GluK2 is sufficient to decrease the levels of aberrantly expressed synaptic GluK2/GluK5 KARs. We also provide evidence that the treatment with AAV9-miRNA#1 is likely to be selective for the *grik2* mRNA as it spares the other KAR subunits as well as major subunits for AMPA, NMDA and GABA_A_ receptors.

Current prospects for gene therapy for focal epilepsy are based on the reduction of neuronal activity, and restoration of the inhibition/excitation balance within the foci to prevent seizures. This is achieved through overexpression of inhibitory peptides such as NPY^58^ or dynorphin^59^ through overexpression of K^+^-channels, such as engineered voltage-gated potassium channel Kv1.1, or through expression of optogenetic and chemogenetic tools^60^ to reduce neuronal excitability^61^. The gene therapy strategy that we advocate is based on a different principle. Instead of dampening the general excitability of neurons by overexpression of an exogenous protein, we seek to selectively downregulate a receptor, GluK2/GluK5, that is aberrantly expressed in a pathological recurrent circuit and promotes chronic recurrent seizures^21^. Thus we directly target a pathological mechanism within the diseased brain. Even moderate suppression of the GluK2 protein expression is sufficient to induce a marked decrease in the frequency of IEDs (both in a mouse model of TLE and in organotypic slices from TLE patients) because removing GluK2 does not block synaptic activity, but it probably prevents the build-up of synaptic depolarization^19^ in response to DGC activity at the start of an epileptic discharge.

In physiological conditions, GluK2/GluK5 KARs play a role in the regulation of developing and mature hippocampal circuits in rodents^53^. The analysis of GluK2 ko mice has provided converging evidence that the overall impact of the genetic manipulation is significant but only subtle at the behavioural level^25, 62^. The constitutive deletion of GluK2 leads to delayed maturation of excitatory synapses in CA3^63^. However, in a similar manner, the selective deletion of GluK2 from developing adult-born DGCs leads to their impaired maturation of synaptic and intrinsic properties, and causes spatial discrimination deficits but no major disruption in a conditioned-fear^64^. Finally, mutations in the *grik2* gene are causative for early childhood development disorders^65^. Hence removing GluK2 in the human hippocampus could potentially cause adverse effects; however, several observations mitigate these concerns: i) GluK2 ko mice only show subtle behavioural deficits^53^, ii) Constitutive deletion of GluK2 in mice does not affect AMPA-mediated EPSCs and prevents presynaptic facilitation at output synapses of the dentate gyrus, thereby leading to decreased spike transfer in physiological conditions^53^; iii) Downregulation of GluK2 in the adult brain bypasses the involvement of GluK2 during postnatal development, iv) Achieving moderate downregulation of GluK2 seems effective in reducing epileptic activity in the mouse models of TLE and in human organotypic slices, and v) hippocampal circuits are highly dysfunctional in TLE patients^66^, hence preventing major recurrent seizures within hippocampal circuits is expected to improve neural circuit activity. To refine our approach, restricting the downregulation of GluK2 to DGCs may also be sufficient to alleviate epileptic activity in TLE.

In conclusion, we employed an innovative pipeline, using a mouse TLE model in parallel to human tissue to deliver proof of concept for our gene therapy strategy for TLE. Overall, our study provides strong evidence that a gene therapy approach selectively targeting GluK2 in the hippocampus is a promising therapeutic strategy.

## Acknowledgments

This work was supported by the Institut National de la Santé et de la Recherche Médicale (INSERM to VC, CB, AP, TM, RK, IK, and FM), the Aix-Marseille University (AMU to VC, CB, AP, TM, RK, IK, and FM), the Centre National de la Recherche Scientifique (CNRS to CM, SD, MM, JV, JR), the Bordeaux University (UB to CM, SD, MM, JV, JR), the AP-HM La Timone Marseille Hospital (to AT, NV, AL, MM, DF-B, ED, ST, RM), the CHU Pellegrin Bordeaux (to GP, CM), the Agence Nationale de la Recherche (ANR 13-BSV4-0012-02 and ANR-18-CE17-0023-01 to CM and VC), Inserm Transfert (to VC), Aquitaine Science Transfert (to CM, JR, MM), REGENXBIO (to AG, AM, JS and OD), and Corlieve Therapeutics (to NP, JG and RP). We thank R. Miles, E. Eugene and C. Le Duigou (ICM, Paris) for their insightful advice on preparing organotypic slices from TLE patients. We also thank E. Pallesi, FJ. Michel and S. Pellegrino-Corby for their technical assistance and heading their respective INMED core facilities, J. Teillon and the Bordeaux Imaging Center (BIC) - a service unit of the CNRS-Inserm and Bordeaux University, member of the national infrastructure France BioImaging-, Y. Rufin from the Biochemistry and Biophysics platform of Bordeaux Neurocampus, E. Verdier and S. Daburon from the Cell Biology Facility of IINS for their technical expertise and providing us respectively rat and mouse primary neurons and the vectorology facility (Vect’UB-TBMCore CNRS UAR3427 - INSERM US005) of Bordeaux University for providing lentiviral particules.

## Author Contributions

CMu and VC contributed to the study concept and design. SD, CB, AP, JG, JV, TM, RK, IK, FM, MM, NP, JR participated in the data acquisition and analysis. DS, FB, GP, CM, SL, AT, NV, AL, MM, DFB, ED, ST, RA, provided experimental human samples. CMu and VC wrote the manuscript with inputs from RP, JS, AM, AG, SB and OD.

## References

1. Kalilani L, Sun X, Pelgrims B, et al. The epidemiology of drug-resistant epilepsy: A systematic review and meta-analysis. Epilepsia 2018;59(12):2179–2193.

2. Tatum WO. Mesial temporal lobe epilepsy. J. Clin. Neurophysiol. Off. Publ. Am. Electroencephalogr. Soc. 2012;29(5):356–365.

3. Black LC, Schefft BK, Howe SR, et al. The effect of seizures on working memory and executive functioning performance. Epilepsy Behav. 2010;17(3):412–419.

4. Kent GP, Schefft BK, Howe SR, et al. The effects of duration of intractable epilepsy on memory function. Epilepsy Behav. EB 2006;9(3):469–477.

5. Monti G, Meletti S. Emotion recognition in temporal lobe epilepsy: A systematic review. Neurosci. Biobehav. Rev. 2015;55:280–293.

6. Jacoby A. Stigma, epilepsy, and quality of life. Epilepsy Behav. EB 2002;3(6S2):10–20.

7. Reinholdson J, Olsson I, Edelvik Tranberg A, Malmgren K. Long-term employment outcomes after epilepsy surgery in childhood. Neurology 2020;94(2):e205–e216.

8. WHO. Epilepsy fact sheet [Internet]. 2022;[cited 2022 Aug 2] Available from: https://www.who.int/news-room/fact-sheets/detail/epilepsy

9. Chen Z, Brodie MJ, Liew D, Kwan P. Treatment Outcomes in Patients With Newly Diagnosed Epilepsy Treated With Established and New Antiepileptic Drugs: A 30-Year Longitudinal Cohort Study. JAMA Neurol. 2018;75(3):279–286.

10. Engel J. THE CURRENT PLACE OF EPILEPSY SURGERY. Curr. Opin. Neurol. 2018;31(2):192–197.

11. Jehi L, Jetté N. Not all that glitters is gold: A guide to surgical trials in epilepsy. Epilepsia Open 2016;1(1–2):22–36.

12. Malmgren K, Edelvik A. Long-term outcomes of surgical treatment for epilepsy in adults with regard to seizures, antiepileptic drug treatment and employment. Seizure 2017;44:217–224.

13. Engel J. What can we do for people with drug-resistant epilepsy? Neurology 2016;87(23):2483– 2489.

14. Represa A, Tremblay E, Ben-Ari Y. Kainate binding sites in the hippocampal mossy fibers: localization and plasticity. Neuroscience 1987;20(3):739–748.

15. Sutula T, Cascino G, Cavazos J, et al. Mossy fiber synaptic reorganization in the epileptic human temporal lobe. Ann. Neurol. 1989;26(3):321–330.

16. Buckmaster PS. Mossy Fiber Sprouting in the Dentate Gyrus [Internet]. In: Noebels JL, Avoli M, Rogawski MA, et al., editors. Jasper’s Basic Mechanisms of the Epilepsies. Bethesda (MD): National Center for Biotechnology Information (US); 2012[cited 2022 Jul 18] Available from: http://www.ncbi.nlm.nih.gov/books/NBK98174/

17. Cavarsan CF, Malheiros J, Hamani C, et al. Is Mossy Fiber Sprouting a Potential Therapeutic Target for Epilepsy? Front. Neurol. 2018;9:1023.

18. Epsztein J, Represa A, Jorquera I, et al. Recurrent mossy fibers establish aberrant kainate receptor-operated synapses on granule cells from epileptic rats. J. Neurosci. Off. J. Soc. Neurosci. 2005;25(36):8229–8239.

19. Artinian J, Peret A, Mircheva Y, et al. Impaired neuronal operation through aberrant intrinsic plasticity in epilepsy: Aberrant Intrinsic Plasticity. Ann. Neurol. 2015;77(4):592–606.

20. Peret A, Christie LA, Ouedraogo DW, et al. Contribution of aberrant GluK2-containing kainate receptors to chronic seizures in temporal lobe epilepsy. Cell Rep. 2014;8(2):347–354.

21. Crépel V, Mulle C. Physiopathology of kainate receptors in epilepsy. Curr. Opin. Pharmacol. 2015;20:83–88.

22. The Assistance Publique Hôpitaux de Marseille’s Biobank [Internet]. [date unknown];[cited 2022 Jul 18] Available from: https://openbioresources.metajnl.com/articles/10.5334/ojb.63/

23. Birmingham A, Anderson E, Sullivan K, et al. A protocol for designing siRNAs with high functionality and specificity. Nat. Protoc. 2007;2(9):2068–2078.

24. Paddison PJ, Cleary M, Silva JM, et al. Cloning of short hairpin RNAs for gene knockdown in mammalian cells. Nat. Methods 2004;1(2):163–167.

25. Mulle C, Sailer A, Pérez-Otaño I, et al. Altered synaptic physiology and reduced susceptibility to kainate-induced seizures in GluR6-deficient mice. Nature 1998;392:601–605.

26. Andersson M, Avaliani N, Svensson A, et al. Optogenetic control of human neurons in organotypic brain cultures. Sci. Rep. 2016;6:24818.

27. Le Duigou C, Savary E, Morin-Brureau M, et al. Imaging pathological activities of human brain tissue in organotypic culture. J. Neurosci. Methods 2018;298:33–44.

28. Eugène E, Cluzeaud F, Cifuentes-Diaz C, et al. An organotypic brain slice preparation from adult patients with temporal lobe epilepsy. J. Neurosci. Methods 2014;235:234–244.

29. Peppi M, Landa M, Sewell WF. Cochlear Kainate Receptors. JARO J. Assoc. Res. Otolaryngol. 2012;13(2):199–208.

30. Epping L, Schroeter CB, Nelke C, et al. Activation of non-classical NMDA receptors by glycine impairs barrier function of brain endothelial cells. Cell. Mol. Life Sci. CMLS 2022;79(9):479.

31. Vigier A, Partouche N, Michel FJ, et al. Substantial outcome improvement using a refined pilocarpine mouse model of temporal lobe epilepsy. Neurobiol. Dis. 2021;161:105547.

32. Cetin A, Komai S, Eliava M, et al. Stereotaxic gene delivery in the rodent brain. Nat. Protoc. 2006;1(6):3166–3173.

33. Kockmann T, Trachsel C, Panse C, et al. Targeted proteomics coming of age - SRM, PRM and DIA performance evaluated from a core facility perspective. Proteomics 2016;16(15–16):2183– 2192.

34. Escher C, Reiter L, MacLean B, et al. Using iRT, a normalized retention time for more targeted measurement of peptides. PROTEOMICS 2012;12(8):1111–1121.

35. Reiter L, Rinner O, Picotti P, et al. mProphet: automated data processing and statistical validation for large-scale SRM experiments. Nat. Methods 2011;8(5):430–435.

36. Müller CJ, Gröticke I, Bankstahl M, Löscher W. Behavioral and cognitive alterations, spontaneous seizures, and neuropathology developing after a pilocarpine-induced status epilepticus in C57BL/6 mice. Exp. Neurol. 2009;219(1):284–297.

37. Twele F, Töllner K, Bankstahl M, Löscher W. The effects of carbamazepine in the intrahippocampal kainate model of temporal lobe epilepsy depend on seizure definition and mouse strain. Epilepsia Open 2016;1(1–2):45–60.

38. Cho K-O, Lybrand ZR, Ito N, et al. Aberrant hippocampal neurogenesis contributes to epilepsy and associated cognitive decline. Nat. Commun. 2015;6:6606.

39. Pinheiro PS, Lanore F, Veran J, et al. Selective block of postsynaptic kainate receptors reveals their function at hippocampal mossy fiber synapses. Cereb. Cortex N. Y. N 1991 2013;23(2):323–331.

40. Dolman NP, More JCA, Alt A, et al. Synthesis and pharmacological characterization of N3-substituted willardiine derivatives: role of the substituent at the 5-position of the uracil ring in the development of highly potent and selective GLUK5 kainate receptor antagonists. J. Med. Chem. 2007;50(7):1558–1570.

41. Perrais D, Pinheiro PS, Jane DE, Mulle C. Antagonism of recombinant and native GluK3-containing kainate receptors. Neuropharmacology 2009;56(1):131–140.

42. Matsuda K, Budisantoso T, Mitakidis N, et al. Transsynaptic Modulation of Kainate Receptor Functions by C1q-like Proteins. Neuron 2016;90(4):752–767.

43. Sabolek HR, Swiercz WB, Lillis KP, et al. A Candidate Mechanism Underlying the Variance of Interictal Spike Propagation. J. Neurosci. 2012;32(9):3009–3021.

44. Bettler B, Mulle C. Review: neurotransmitter receptors. II. AMPA and kainate receptors. Neuropharmacology 1995;34(2):123–139.

45. Sills GJ, Rogawski MA. Mechanisms of action of currently used antiseizure drugs. Neuropharmacology 2020;168:107966.

46. Kaminski RM, Banerjee M, Rogawski MA. Topiramate selectively protects against seizures induced by ATPA, a GluR5 kainate receptor agonist. Neuropharmacology 2004;46(8):1097– 1104.

47. Taniguchi S, Stolz JR, Swanson GT. The antiseizure drug perampanel is a subunit-selective negative allosteric modulator of kainate receptors. J. Neurosci. Off. J. Soc. Neurosci. 2022;JN-RM-2397-21.

48. Gibbs III JW, Sombati S, DeLorenzo RJ, Coulter DA. Cellular Actions of Topiramate: Blockade of Kainate-Evoked Inward Currents in Cultured Hippocampal Neurons. Epilepsia 2000;41(s1):10– 16.

49. Hanada T. The discovery and development of perampanel for the treatment of epilepsy. Expert Opin. Drug Discov. 2014;9(4):449–458.

50. Hesdorffer DC, Kanner AM. The FDA alert on suicidality and antiepileptic drugs: Fire or false alarm? Epilepsia 2009;50(5):978–986.

51. Ettinger AB, LoPresti A, Yang H, et al. Psychiatric and behavioral adverse events in randomized clinical studies of the noncompetitive AMPA receptor antagonist perampanel. Epilepsia 2015;56(8):1252–1263.

52. Wenthold RJ, Trumpy VA, Zhu WS, Petralia RS. Biochemical and assembly properties of GluR6 and KA2, two members of the kainate receptor family, determined with subunit-specific antibodies. J. Biol. Chem. 1994;269(2):1332–1339.

53. Mulle C, Crépel V. Regulation and dysregulation of neuronal circuits by KARs. Neuropharmacology 2021;197:108699.

54. Kantor B, Bailey RM, Wimberly K, et al. Methods for gene transfer to the central nervous system. Adv. Genet. 2014;87:125–197.

55. Artinian J, Peret A, Marti G, et al. Synaptic Kainate Receptors in Interplay with INaP Shift the Sparse Firing of Dentate Granule Cells to a Sustained Rhythmic Mode in Temporal Lobe Epilepsy. J. Neurosci. 2011;31(30):10811–10818.

56. Ruiz A, Sachidhanandam S, Utvik JK, et al. Distinct subunits in heteromeric kainate receptors mediate ionotropic and metabotropic function at hippocampal mossy fiber synapses. J. Neurosci. Off. J. Soc. Neurosci. 2005;25(50):11710–11718.

57. Christensen JK, Paternain AV, Selak S, et al. A mosaic of functional kainate receptors in hippocampal interneurons. J. Neurosci. Off. J. Soc. Neurosci. 2004;24(41):8986–8993.

58. Woldbye DPD, Angehagen M, Gøtzsche CR, et al. Adeno-associated viral vector-induced overexpression of neuropeptide Y Y2 receptors in the hippocampus suppresses seizures. Brain J. Neurol. 2010;133(9):2778–2788.

59. Agostinho AS, Mietzsch M, Zangrandi L, et al. Dynorphin-based “release on demand” gene therapy for drug-resistant temporal lobe epilepsy. EMBO Mol. Med. 2019;11(10):e9963.

60. Walker MC, Kullmann DM. Optogenetic and chemogenetic therapies for epilepsy. Neuropharmacology 2020;168:107751.

61. Snowball A, Chabrol E, Wykes RC, et al. Epilepsy Gene Therapy Using an Engineered Potassium Channel. J. Neurosci. Off. J. Soc. Neurosci. 2019;39(16):3159–3169.

62. Micheau J, Vimeney A, Normand E, et al. Impaired hippocampus-dependent spatial flexibility and sociability represent autism-like phenotypes in GluK2 mice. Hippocampus 2014;24(9):1059–1069.

63. Lanore F, Labrousse VF, Szabo Z, et al. Deficits in Morphofunctional Maturation of Hippocampal Mossy Fiber Synapses in a Mouse Model of Intellectual Disability. J. Neurosci. 2012;32(49):17882–17893.

64. Zhu Y, Armstrong JN, Contractor A. Kainate receptors regulate the functional properties of young adult-born dentate granule cells. Cell Rep. 2021;36(12):109751.

65. Stolz JR, Foote KM, Veenstra-Knol HE, et al. Clustered mutations in the GRIK2 kainate receptor subunit gene underlie diverse neurodevelopmental disorders. Am. J. Hum. Genet. 2021;108(9):1692–1709.

66. Gabriel S, Njunting M, Pomper JK, et al. Stimulus and potassium-induced epileptiform activity in the human dentate gyrus from patients with and without hippocampal sclerosis. J. Neurosci. Off. J. Soc. Neurosci. 2004;24(46):10416–10430.

